# Sox4 in Treg Cells Suppresses IL-10 Production via c-Maf Degradation and Exacerbates Type 2 Inflammation

**DOI:** 10.1101/2025.11.04.686669

**Authors:** Yuki Hayashi, Kensuke Suga, Akira Suto, Arifumi Iwata, Junichi Ishikawa, Kazuya Abe, Takahiro Kageyama, Takashi Ito, Shigeru Tanaka, Kotaro Suzuki, Masakatsu Yamashita, Véronique Lefebvre, Hiroshi Nakajima

**Affiliations:** Department of Allergy and Clinical Immunology, Graduate School of Medicine, Chiba University, Chiba, Japan; Institute for Advanced Academic Research, Chiba University, Chiba, Japan; Chiba University Synergy Institute for Futuristic Mucosal Vaccine Research and Development (cSIMVa), Chiba University, Chiba, Japan; Department of Infections and Host Defenses, Ehime University Graduate School of Medicine, Ehime, Japan; Department of Immunology, Ehime University Graduate School of Medicine, Ehime, Japan; Department of Surgery, Division of Orthopaedic Surgery, Children’s Hospital of Philadelphia, Pennsylvania, USA

## Abstract

The anti-inflammatory cytokine interleukin-10 (IL-10) produced by regulatory T (Treg) cells plays a crucial role in regulating allergic airway inflammation. However, the mechanisms governing IL-10 production in Treg cells remain unclear. In this study, we first identified *Sox4* as a gene whose expression is down-regulated in IL-10-producing Treg cells. Comprehensive gene expression profiling of Sox4-deficient Treg cells and Sox4-overexpressing Treg cells revealed specific downregulation of *Il10* and *Ctla4* by Sox4. Sox4-deficient Treg cells exhibited increased IL-10 production and CTLA-4 expression both in vitro and in vivo and strongly suppressed Th2 cell proliferation compared with Sox4-expressing Treg cells. In a house dust mite (HDM)-induced asthma model, Treg-specific Sox4-deficient (Sox4-cKO) mice exhibited IL-10-dependent reductions in eosinophilic inflammation and Th2 cell infiltration in the airways. A comparison of Sox4-cKO and wild-type Treg cells in the same inflamed environment in mixed bone marrow chimeric mice with HDM-induced asthma revealed that Sox4-mediated suppression of c-Maf is cell-intrinsic. In addition, overexpression of Sox4 in Treg cells repressed the activation of c-Maf-regulated chromatin regions, including those around the *Il10* and *Ctla4* loci. Moreover, Sox4-induced suppression of IL-10 and CTLA-4 in Treg cells was rescued by forced expression of c-Maf. While Sox4 did not significantly affect *Maf* mRNA levels, Sox4 degraded c-Maf proteins in Treg cells through its 33 C-terminal residues. Via a domain distinct from its C-terminus, Sox4 physically interacted with c-Maf, subsequently facilitating its degradation through a ubiquitin-proteasome pathway. These findings elucidate a mechanism by which Sox4 influences the asthmatic responses by regulating the c-Maf/IL-10 axis in lung Treg cells.

## INTRODUCTION

Allergic asthma is triggered by sensitization to inhaled antigens, such as house dust mites (HDM), and is characterized by chronic eosinophilic and type 2 inflammation^1, 2^. This condition is mainly mediated by T helper 2 (Th2) cells, whose master regulator is GATA3. Conversely, regulatory T (Treg) cells, whose master regulator is Foxp3, act to suppress the allergic responses^3^. The differentiation of Th2 cells from naïve CD4^+^ T cells occurs upon stimulation by antigen-presenting cells in the presence of IL-4. Once differentiated, Th2 cells produce cytokines such as IL-5 and IL-13, which play crucial roles in the pathogenesis of allergic asthma. Specifically, IL-5 promotes eosinophil infiltration into the airways, while IL-13 contributes to airway hyperresponsiveness.

Interleukin-10 (IL-10), a potent anti-inflammatory cytokine, suppresses allergic airway inflammation^4, 5, 6^. Initially, the regulatory effects of IL-10 on T cell-mediated responses were thought to be indirect, primarily occurring through antigen-presenting cells^7^. However, recent studies have revealed that IL-10 can directly influence both IL-17-producing CD4^+^ T (Th17) cells and Treg cells, as demonstrated in murine colitis models^8, 9^. In the context of allergic inflammation, recent research has shown that IL-10 exerts its effects by directly inhibiting Th2 cell differentiation and survival^10^.

Regarding IL-10-producing T cells, in vitro studies have demonstrated that efficient IL-10 production from helper T cells requires stimulation with vitamin D3 and dexamethasone^11^. In vivo disease models, including malaria infection and allergic asthma, have shown that Th1 and Th2 cells can produce IL-10. This production is mediated through the expression of the transcription factor c-Maf, highlighting a common molecular pathway across different T cell subsets for IL-10 production^12^. On the other hand, recent in vitro studies have revealed the cytokine milieu that promotes IL-10 production in Treg cells. Zhou et al. have shown that IL-10-producing Treg cells are synergistically induced by the combined action of IL-2 and IL-4^13^. Given that IL-2 is essential for Treg cell survival and that IL-4 is a critical cytokine in Th2 cell differentiation, these findings indicate the significance of IL-10-producing Treg cells in suppressing inflammation induced by Th2 cells. Indeed, they have also shown that combination therapy of IL-2 and IL-4 ameliorates the HDM-induced murine asthma model through IL-10 production.

Despite the capacity for IL-10 production by various T cell subsets, IL-10-producing Treg cells play a particularly crucial role in suppressing allergic eosinophilic inflammation. This notion is evidenced by increased eosinophilic airway inflammation in Treg-specific IL-10-deficient mice^14^, contrasted with decreased eosinophilic inflammation in T cell-specific IL-10-deficient mice^15^. Therefore, elucidating the mechanisms governing IL-10 production in Treg cells will be critical for developing novel strategies to resolve allergic eosinophilic inflammation.

In this study, to further investigate IL-10-producing Treg cells, we first re-analyzed public RNA-sequencing datasets for IL-10-producing T cells. We found that *Sox4* was the only gene for a transcription factor to be down-regulated in IL-10-producing Treg cells, suggesting its potential role in regulating IL-10 production in Treg cells. Sox4 is a member of the SOXC group within the SOX family. This group includes Sox4, Sox11, and Sox12. The SoxC family members contain an HMG-type DNA-binding domain and a 33-residue-long C-terminal region that acts as a transactivation domain and a target site to induce proteasomal degradation^16, 17^. Regarding the role of SoxC family proteins in Treg cells, although Sox12 induces Foxp3 expression in peripherally induced Treg cells upon TCR stimulation, such stimulation decreases *Sox4* expression but increases *Sox12* expression in Treg cells, suggesting that these two SoxC proteins play distinct roles in Treg cell function^18^. We demonstrate here that Sox4 expressed in Treg cells suppresses IL-10 production by decreasing c-Maf proteins and that Sox4 in Treg cells enhances HDM-induced eosinophilic inflammation.

## RESULTS

### *Sox4* is downregulated in IL-10-producing Treg cells

To identify the genes crucial for IL-10-producing Treg cells, we first re-analyzed a public RNA-sequencing dataset for IL-10-producing Treg cells (IL-10^+^ Treg) and IL-10-producing CD4^+^ T cells that were induced with vitamin D3 and dexamethasone (IL-10^+^ Tconv (VitD3+Dex))^12^. By comparing pairs of IL-10^+^ Treg and IL-10 non-producing Treg cells (IL-10^−^ Treg) (pair 1) and of IL-10^+^ Treg and IL-10^+^ Tconv (VitD3+Dex) (pair 2), we extracted 44 differentially expressed genes (DEG) from pair 1 and 535 DEG from pair 2 and found that 36 genes were shared in the two pairs (Figure 1A). Among them, we found only three encoding transcription factors: *Sox4* was downregulated, and *Rorc* and *Arnt2* were upregulated in IL-10-producing Treg cells.

**Figure 1.**
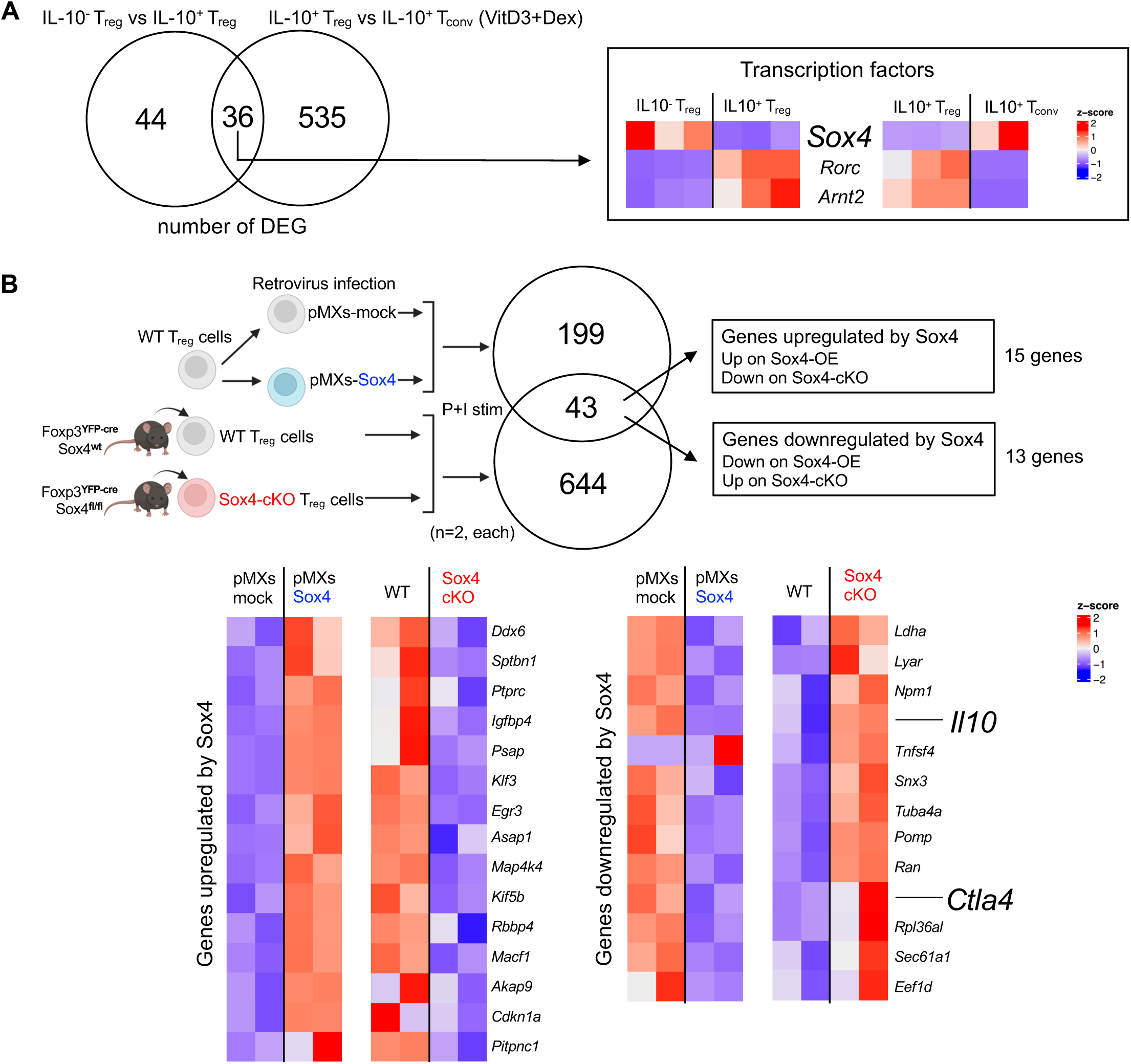
*Sox4* is downregulated in IL-10-producing Treg cells. (**A**) Re-analysis of IL-10 producing cells: RNA-seq data of Foxp3RFP^+^ IL-10GFP^+^ CD4^+^ T cells (IL-10^+^ Treg) and Foxp3RFP^+^ IL-10GFP^-^ CD4^+^ T cells (IL-10^-^ Treg) directly sorted from Foxp3^RFP^ IL-10^GFP^ mice (n = 3, each) were re-analyzed. IL-10^+^ CD4^+^ T cells induced from naïve CD4^+^ T cells by stimulating anti-CD3 antibody (Ab) and anti-CD28 antibody in the presence of vitamin D3 and dexamethasone plus anti-IL-4 Ab, anti-IFN-γ Ab, and anti-IL-12 Ab for 21 days (IL-10^+^ Tconv (VitD3+Dex)) (n = 2) were also re-analyzed. By comparing pairs of IL-10^+^ Treg and IL-10^−^ Treg (pair 1) and of IL-10^+^ Treg and IL-10^+^ Tconv (VitD3+Dex) (pair 2), 44 DEG from pair 1 and 535 DEG from pair 2 were detected. Among 36 shared genes, three transcription factors (*Sox4*, *Rorc*, *Arnt2*) were detected. (**B**) For DEG on Sox4 overexpressing (Sox4-OE) Treg cells, Treg cells purified from Foxp3^YFP-cre^ mice (WT Treg cells) were cultured in IL-10^+^Treg-polarizing conditions and infected with retrovirus of pMXs-IRES-NGFR (pMXs-mock) or pMXs-Sox4-IRES-NGFR (pMXs-Sox4). Infected cells were sorted 3 days after the infection, stimulated with PMA and ionomycin (P+I), and subjected to RNA-sequence analysis. For DEG on Sox4 conditional knockout (Sox4-cKO) Treg cells, Treg cells purified from Foxp3^YFP-cre^ mice (WT) or Foxp3^YFP-cre^ Sox4^f/f^ mice (Sox4-cKO) were stimulated with P+I and subjected to RNA-sequence analysis (n=2, each). Among genes shared by the two comparisons, fifteen genes were upregulated, and thirteen genes were downregulated by Sox4.

Recent studies have shown that the proliferation of IL-10-producing Treg cells is facilitated by the collaboration of IL-4, a critical factor in Th2 differentiation, and IL-2, a critical factor for Treg survival^13^. Given that Sox4 has been shown to suppress Th2 cell function,^19^ it is possible that Sox4 affects the function of IL-10-producing Treg cells. Accordingly, we next searched for the genes whose expression is regulated by Sox4 in Treg cells. In this experiment, Treg cells were isolated from the spleen and lymph nodes of Foxp3^YFP-cre/YFP-cre^ mice and Foxp3^YFP-cre/YFP-cre^ Sox4^f/f^ mice (WT Treg cells and Sox4-cKO Treg cells, respectively, n = 2, each). In another set of experiments, sorted WT Treg cells were infected with pMXs-Sox4-IRES-NGFR retrovirus (pMXs-Sox4) or control pMXs-IRES-NGFR retrovirus (pMXs-mock), and the infected cells were isolated by the expression of NGFR (n = 2, each) (Figure 1B). These cells were stimulated with PMA and ionomycin (P+I) and subjected to RNA-sequencing. By comparing pairs of WT Treg cells and Sox4-cKO Treg cells and of pMXs-Sox4 and pMXs-mock, we extracted the shared DEG between the pairs and found that 13 genes were downregulated, while 15 genes were upregulated by Sox4. Among them, *Il10* and *Ctla4* were downregulated by Sox4.

### Sox4-cKO Treg cells increase the expression of IL-10 and CTLA-4 and suppress Th2 cells

Next, we investigated whether Sox4 regulates IL-10 and CTLA-4 at protein levels. We found that IL-10 production and CTLA-4 expression were diminished in WT Treg cells infected with pMXs-Sox4 relative to those infected with pMXs–-mock (Figure 2A, left panels). IL-10 production and CTLA-4 expression were consistently elevated in Sox4-cKO Treg cells compared to WT Treg cells (Figure 2A, right panels). These results indicate that Sox4 suppresses IL-10 production and CTLA-4 expression in Treg cells. We also found that Sox4-cKO Treg cells showed increased suppression activity compared to WT Treg cells in the Treg suppression assay (Figure 2B). Since soluble factors, such as IL-10, are less critical for the Treg suppression assay in vitro^20^, Sox4-mediated downregulation of *Ctla4* (Figure 1B) may impact this assay. To further determine whether the enhanced suppressive activity observed in Sox4*-*cKO Treg cells was mediated by CTLA-4, we treated the cells with an anti-CTLA-4 antibody or an isotype control antibody (Figure 2C). CTLA-4 neutralization abolished the difference in suppressive activity between Sox4*-*cKO and WT Treg cells, indicating that Sox4 downregulates CTLA-4-dependent suppressive function.

**Figure 2.**
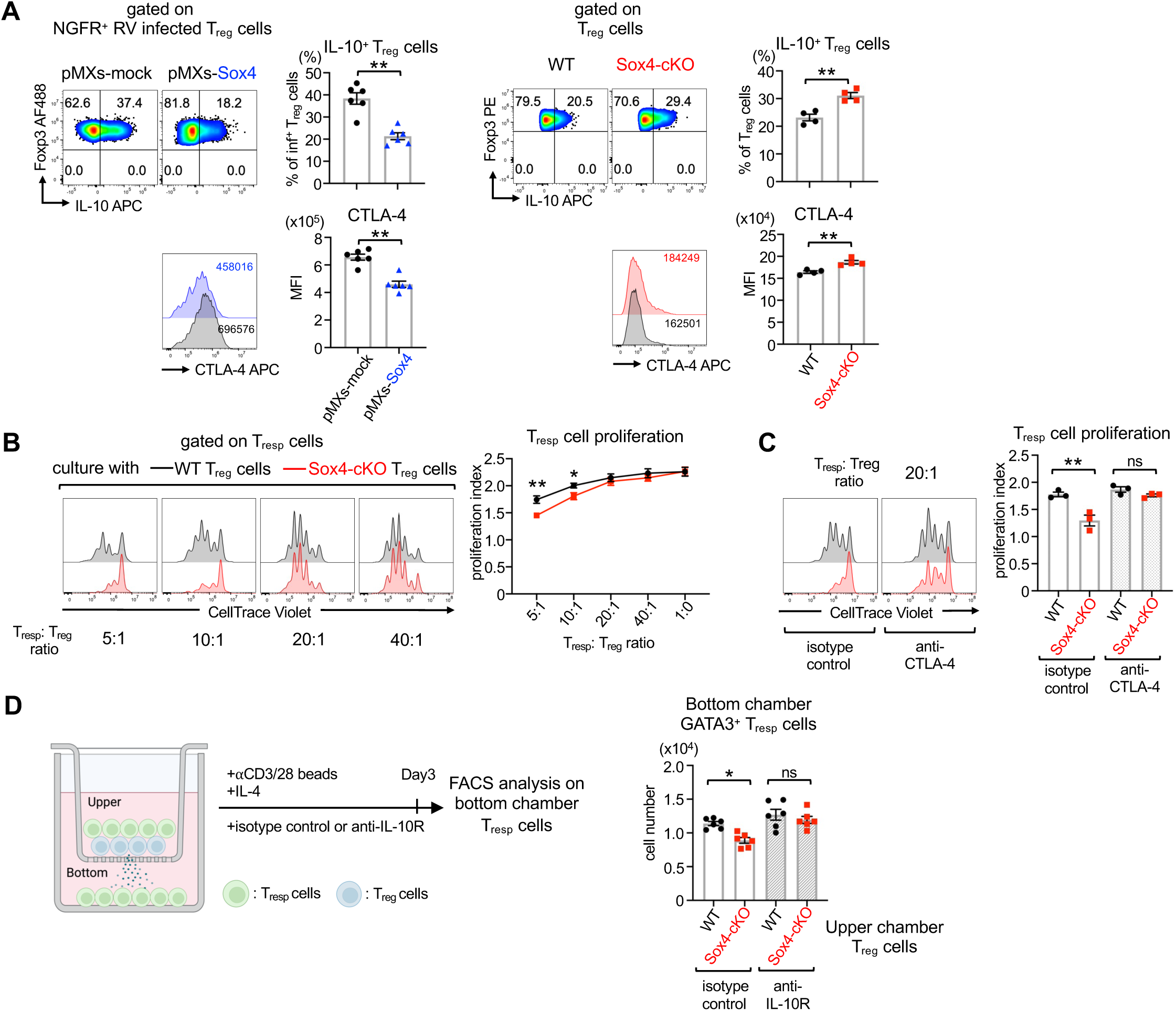
Sox4-cKO Treg cells increase IL-10 and CTLA-4 and suppress Th2 cells. (**A**) Left panels: Treg cells purified from Foxp3^YFP-cre^ mice (WT Treg cells) were cultured and infected with retrovirus as described in Figure 1B. Representative flow cytometry profiles and the frequency of IL-10^+^ Treg cells and the MFI of CTLA-4 on Foxp3^+^ NGFR^+^ infected cells are shown (n = 6 each). Right panels: WT and Sox4-cKO Treg cells were purified and stimulated as described in Figure 1B. Representative flow cytometry profiles and the frequency of IL-10^+^ Treg cells and the MFI of CTLA-4 on Foxp3^+^ cells are shown (n = 4 each). Unpaired t-test, **P<0.01. (**B**) WT or Sox4-cKO Treg cells were co-cultured with irradiated splenocytes and CellTrace Violet-labeled naïve CD45.1^+^ CD4^+^ T cells (Tresp) at the indicated ratio in the presence of anti-CD3 Abs. Representative histograms of CellTrace Violet on CD45.1^+^ CD4^+^ T cells and means ± SEM of the proliferation index of responder cells are shown. n = 6, Two-way ANOVA followed by Tukey’s test. *p<0.05. **p<0.01. (**C**) Representative histograms of CellTrace Violet on CD45.1^+^ CD4^+^ T cells and means ± SEM of the proliferation index of responder cells in the presence of anti-CTLA-4 or isotype control antibody. n = 3, Two-way ANOVA followed by Tukey’s test. **p<0.01. ns: not significant. (**D**) Left panels: Schema of the transwell experiment. Naïve CD4^+^ T cells (Tresp), WT or Sox4-cKO Treg cells, and anti-CD3/CD28 beads were in the upper chamber, and Tresp and anti-CD3/CD28 beads were in the bottom chamber. Medium was supplemented with IL-4 and, where indicated, with anti-IL-10R antibodies. Right panels: The number of GATA3^+^ Th2 cells in the bottom chamber on day 3. Two-way ANOVA followed by Tukey’s test. n = 6, each. *p<0.05. ns: not significant.

We next examined whether soluble factors produced by Sox4-cKO Treg cells influence the differentiation of Th2 cells, as IL-10 has been shown to suppress Th2 cell differentiation directly^10^. As shown in Figure 2D, Sox4-cKO Treg cells are more suppressive to Th2 cells than WT Treg cells in the transwell experiment. Furthermore, blocking the IL-10 receptor canceled the difference in the effect on the proliferation of Th2 cells between Sox4-cKO Treg cells and WT Treg cells. These results indicate that deficiency of Sox4 in Treg cells results in the enhancement of IL-10 production and CTLA-4 expression, as well as suppressive activity to Th2 cells in vitro.

### Sox4 deficiency in Treg cells ameliorates HDM-induced type 2 inflammation

To determine the role of Sox4 in Treg cells in vivo in type-2 inflammation, we analyzed HDM-induced asthma models in Treg-specific Sox4-deficient mice [Foxp3^YFP-cre/YFP-cre^ Sox4^f/f^(Sox4-cKO)] mice and used Sox4-heterozygous mice [Foxp3^YFP-cre/YFP-cre^ Sox4^f/wild-type^ (Sox4-cHet)] as controls (Figure 3A). Sox4-cKO mice showed significantly lower numbers of eosinophils and CD101^+^ inflammatory eosinophils in the lungs (Figure 3B). In addition, IL-5-producing and IL-13-producing Th2 cells were significantly decreased in number in the lungs of Sox4-cKO mice (Figure 3C). Histological analyses revealed that lung inflammation and goblet cell hyperplasia were significantly reduced in Sox4-cKO mice compared with Sox4-cHet mice (Figure 3D). These results suggest that a deficiency of Sox4 in Treg cells decreases Th2 cell responses and eosinophilic inflammation in the airways.

**Figure 3.**
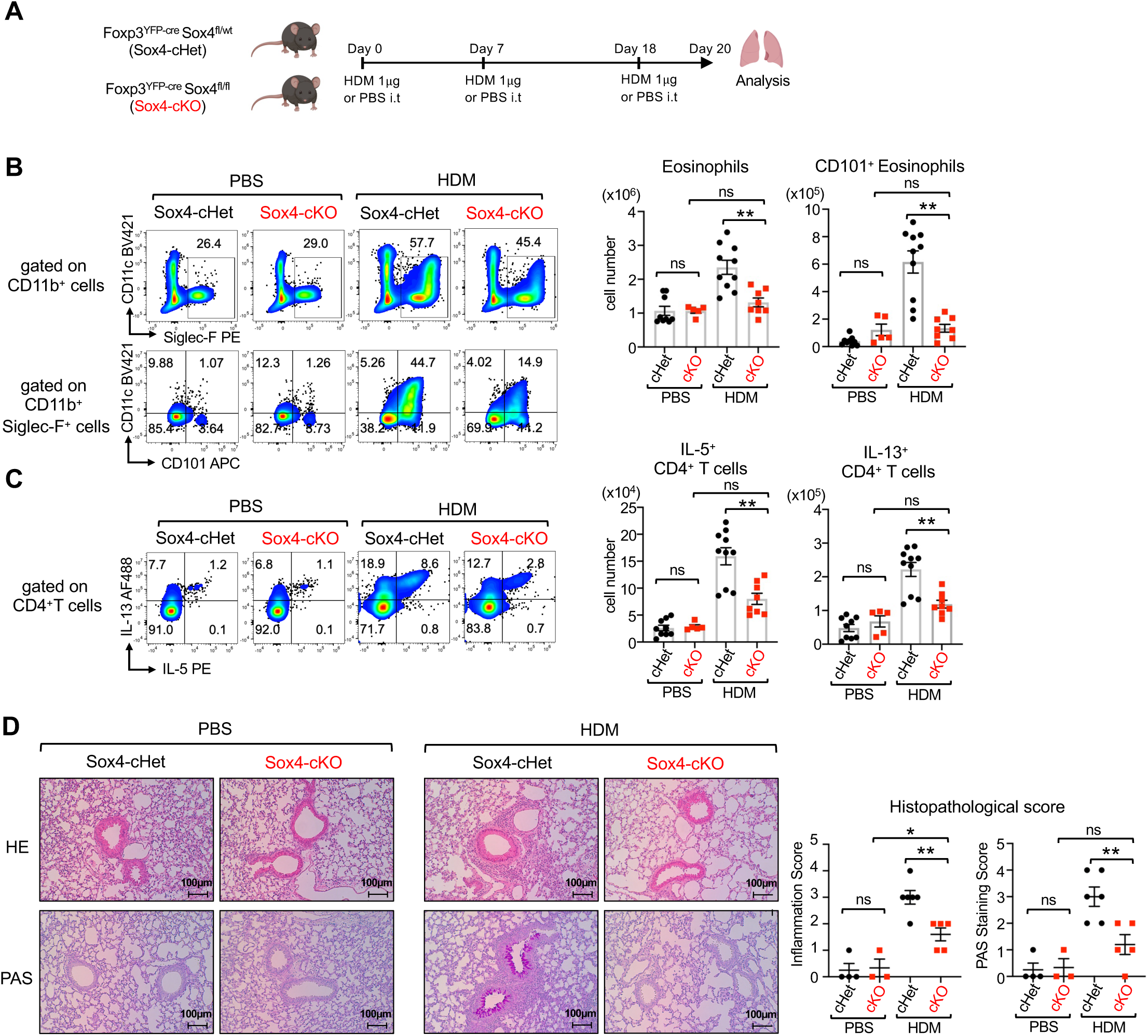
Sox4 deficiency in Treg cells ameliorates HDM-induced type 2 inflammation. (**A**) Shema of HDM-induced asthma model: Foxp3^YFP-cre/YFP-cre^ Sox4^f/f^ (Sox4-cKO) mice and Foxp3^YFP-cre/YFP-cre^ Sox4^f/wt^ (Sox4-cHet) mice were intratracheally sensitized and challenged with HDM (1 μg) or PBS (as control) as described in the Star Methods. (**B**) Representative flow cytometry profiles of CD11c vs. Siglec-F gated on CD45^+^ CD11b^+^ cells and CD11c vs. CD101 gated on CD45^+^ CD11b^+^ Siglec-F^+^ cells in the lung (left panels) and means ± SEM of the numbers of indicated cells in the lung (right panels). (**C**) Representative flow cytometry profiles of IL-5 vs. IL-13 of CD4^+^ T cells in the lung (left panels) and means ± SEM of the numbers of indicated cells in the lung (right panels). n = 9 for Sox4-cHet PBS, n = 5 for Sox4-cKO PBS, n = 10 for Sox4-cHet HDM, and n = 8 for Sox4-cKO HDM. One-way ANOVA. **P<0.01. ns: not significant. (**D**) Representative histological images and histological scores of lung sections. Scale bar = 100 µm. n = 4 for Sox4-cHet PBS, n = 3 for Sox4-cKO PBS, n = 6 for Sox4-cHet HDM, and n = 5 for Sox4-cKO HDM. One-way ANOVA. *p<0.05. **P<0.01. ns: not significant.

### IL-10–dependent amelioration of HDM-induced type 2 inflammation in Sox4-cKO mice

To determine whether the reduced type 2 inflammation observed in Sox4-cKO mice compared with Sox4-cHet mice depends on IL-10, we administered IL-10–neutralizing antibodies in an HDM-induced asthma model and evaluated the impact on inflammatory responses (Figure 4A). Treatment with anti-IL-10 receptor (IL-10R) antibodies before the allergen challenge largely abolished the differences in total eosinophils and CD101□ inflammatory eosinophils in the lungs (Figure 4B). In addition, the differences in IL-5*-* and IL-13*-*producing Th2 cells between Sox4-cKO and Sox4-cHet mice were also largely diminished (Figure 4C). These results indicate that Sox4*-*deficient Treg cells suppress allergic airway inflammation in an IL-10*-*dependent manner in vivo.

**Figure 4.**
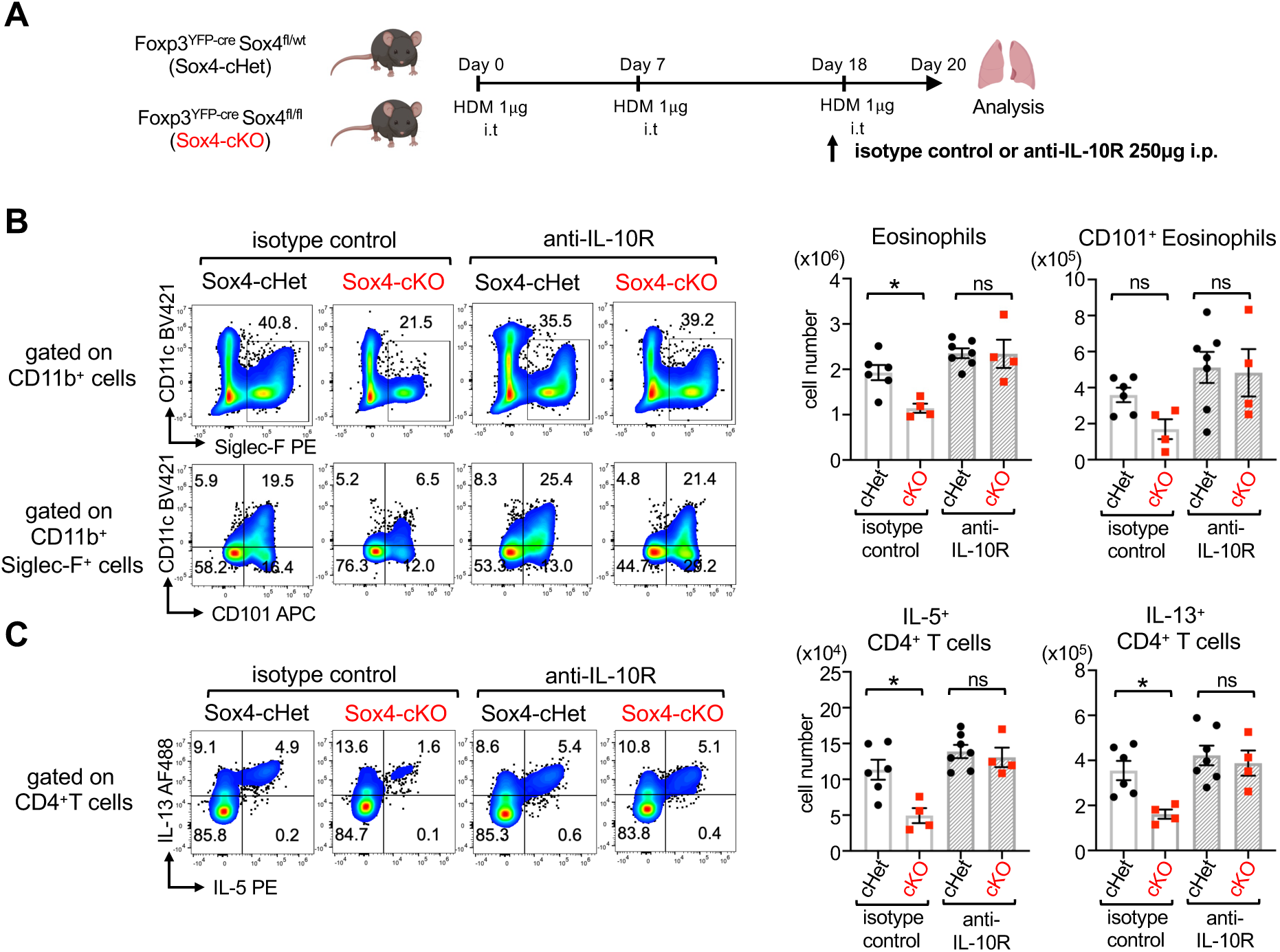
IL-10–dependent amelioration of HDM-induced type 2 inflammation in Sox4-cKO mice. **(A)** Foxp3^YFP-cre/YFP-cre^ Sox4^f/f^ (Sox4-cKO) mice and Foxp3^YFP-cre/YFP-cre^ Sox4^f/wt^ (Sox4-cHet) mice were sensitized and challenged with HDM or PBS (control), and lung cells were analyzed as described in Figure 3A. Anti-IL-10R antibody or isotype control was intraperitoneally administered prior to the challenge. (**B**) Representative flow cytometry profiles of CD11c vs. Siglec-F gated on CD45^+^ CD11b^+^ cells and CD11c vs. CD101 gated on CD45^+^ CD11b^+^ Siglec-F^+^ cells in the lung (left panels) and means ± SEM of the numbers of indicated cells in the lung (right panels). (**C**) Representative flow cytometry profiles of IL-5 vs. IL-13 of CD4^+^ T cells in the lung (left panels) and means ± SEM of the numbers of indicated cells in the lung (right panels). n = 6 for Sox4-cHet isotype control, n = 4 for Sox4-cKO isotype control and anti-IL-10R, n = 7 for Sox4-cHet anti-IL-10R. Two-way ANOVA. *P<0.05. **P<0.01. ns: not significant.

### Sox4-cKO Treg cells increase the expression of GATA3, c-Maf, CTLA-4, and IL-10 under type 2 inflammation

To compare the features of Sox4-deficient Treg cells and Sox4-expressing Treg cells in the same environment, we analyzed HDM-induced asthma models in Foxp3^YFP-cre/wild-type^ Sox4^f/f^ female mice. In this model, since the Foxp3 gene is located on chromosome X and since X chromosome inactivation can occur on either allele, 50% of Treg cells express the Foxp3^YFP-cre^ marker, while the remaining Treg cells do not express Foxp3^YFP-cre^ in Foxp3^YFP-cre/wild-type^ female mice^21^. This model allowed a direct comparison of Foxp3^YFP-cre^ Sox4^f/f^ (Sox4-cKO) Treg cells and Sox4^f/f^ (Sox4-cWT) Treg cells in the same lungs (Figure 5A). We found that IL-10 production and CTLA-4 expression significantly increased in Sox4-cKO Treg cells compared to those in Sox4-cWT Treg cells in the lungs in the HDM-induced asthma model (Figure 5B). We also found that the mean fluorescent intensity (MFI) of GATA3 and c-Maf, but not RORγt, was significantly increased in Sox4-cKO Treg as compared to those in Sox4-cWT Treg cells in the lungs of the HDM-induced asthma model (Figure 5B), although the genes for these transcription factors were not detected as Sox4-regulated genes in the experiment shown in Figure 1B.

**Figure 5.**
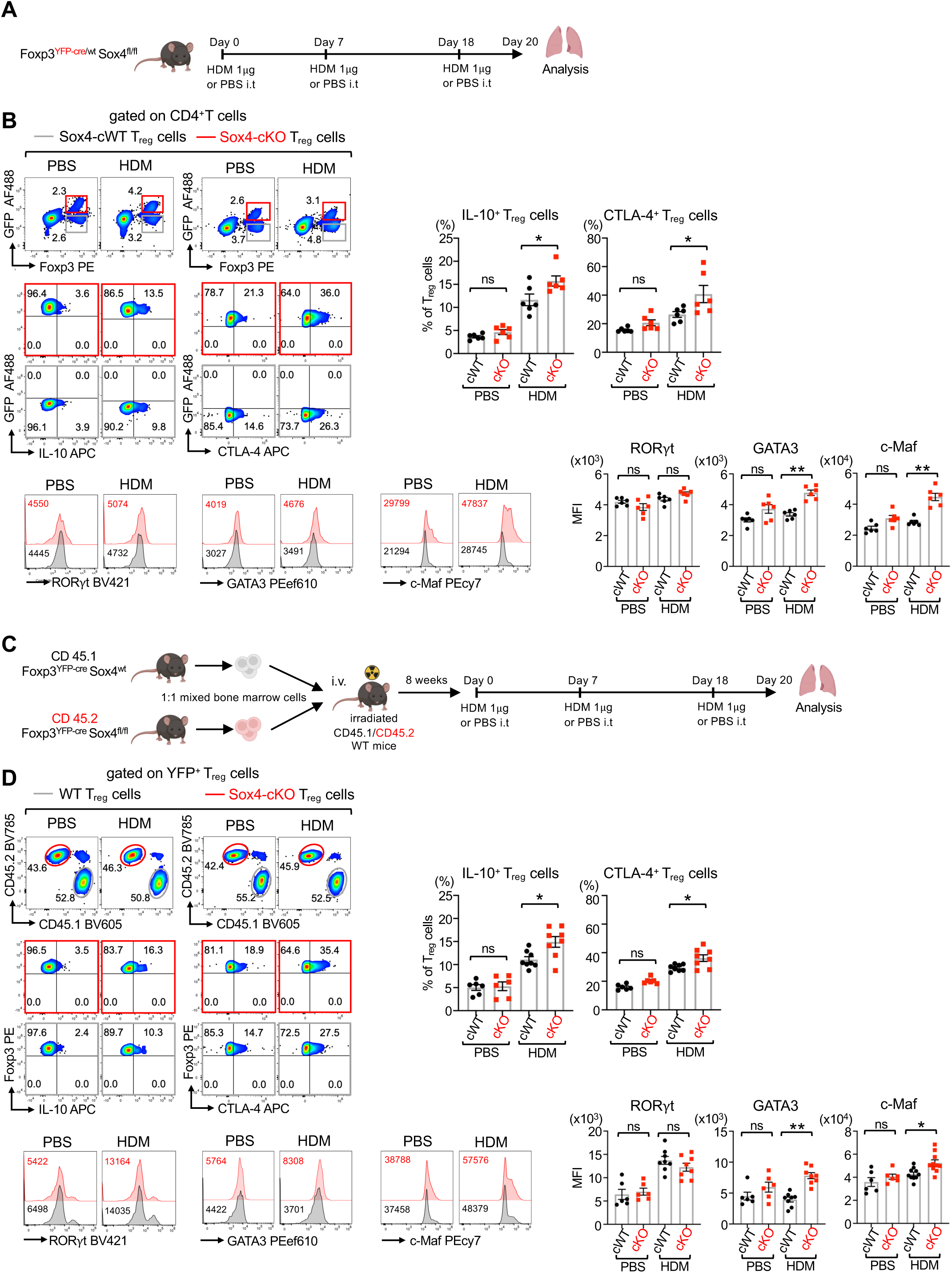
Sox4-cKO Treg cells increase GATA3, c-Maf, CTLA-4, and IL-10 in vivo. (**A**) Foxp3^YFP-cre/wild-type^ Sox4^f/f^ female mice were sensitized and challenged with HDM or PBS (control), and lung cells were analyzed as described in Figure 3A. Intracellular cytokines were analyzed after stimulation with P+I. CTLA-4 and transcription factors were analyzed without in vitro stimulation. (**B**) Upper left panels: Representative flow cytometry profiles of Foxp3 vs. YFP (detected by anti-GFP AF488) gated on CD4^+^ T cells and IL-10 vs. YFP and CTLA-4 vs. YFP gated on either Sox4-cKO Treg cells or Sox4-cWT Treg cells. Sox4-cWT Treg cells were surrounded by gray lines, and Sox4-cKO Treg cells were surrounded by red lines. Upper right panels: Frequencies of IL-10^+^ Treg cells and CTLA-4^+^ Treg cells among the indicated Treg cells. Lower panels: Representative histograms of indicated transcription factors in Sox4-cWT (gray line) and Sox4-cKO (red line) Treg cells and the MFI of indicated transcription factors in Sox4-cWT and Sox4-cKO Treg cells. One-way ANOVA followed by Tukey’s test. n = 6 each. *P<0.05. **P<0.01. ns: not significant. (**C**) Schema of mixed bone marrow chimeric (mBMC) mice: Bone marrow cells from CD45.1^+^ Foxp3^YFP-cre^ (WT) and CD45.2^+^ Foxp3^YFP-cre^ Sox4^f/f^ (Sox4-cKO) mice were mixed at a 1:1 ratio and intravenously injected into lethally irradiated CD45.1^+^/CD45.2^+^ WT mice. Eight weeks after BM cell transfer, mBMC mice were sensitized and challenged with HDM and analyzed as described in Figure 4A. (**D**) Upper left panels: Representative flow cytometry profiles of CD45.1 vs. CD45.2 gated on YFP^+^ Treg cells and IL-10 vs. Foxp3 and CTLA-4 vs. Foxp3 gated on either Sox4-cKO Treg cells (surrounded by red lines) or WT Treg cells (surrounded by grey lines). Upper right panels: Frequencies of IL-10^+^ Treg cells and CTLA-4^+^ Treg cells among the indicated Treg cells. Lower panels: Representative histograms of indicated transcription factors in WT (gray line) and Sox4-cKO (red line) Treg cells and the MFI of indicated transcription factors. n = 8 for HDM and n = 6 for PBS. One-way ANOVA *P<0.05. **P<0.01. ns: not significant.

To further investigate the difference between Sox4-cKO Treg cells and WT Treg cells under the same environmental conditions, we generated mixed bone marrow chimeric (mBMC) mice in which a mixture of CD45.1^+^Foxp3^YFP-cre/YFP-cre^ and CD45.2^+^Foxp3^YFP-cre/YFP-cre^ Sox4^f/f^ bone marrow cells was injected intravenously into lethally irradiated adult CD45.1^+^/CD45.2^+^ mice. Eight weeks after transplantation, the mBMC mice were subjected to the HDM-induced asthma model (Figure 5C). In the lungs, the frequencies of IL-10-producing and CTLA-4^+^ Treg cells were significantly higher in Sox4-cKO Treg cells compared with WT Treg cells (Figure 5D). Additionally, MFI of GATA3 and c-Maf markedly increased in Sox4-cKO Treg cells, whereas RORγt expression remained unchanged. In contrast, no significant difference in these markers was observed between Sox4-cKO and WT Treg cells in the draining mediastinal lymph nodes, either in Foxp3^YFP-cre/wild-type^ Sox4^f/f^ female mice or in mBMC mice (Supplementary Figure 1). Collectively, these findings indicate that the type 2 inflammatory milieu in the lung is required for Sox4-dependent suppression of IL-10 production in Treg cells. Taken together, Sox4 deficiency in Treg cells enhances c-Maf and GATA3 expression, as well as IL-10 production and CTLA-4 expression, in a pulmonary type 2 inflammatory environment.

### Sox4 decreases IL-10 and CTLA-4 expression via suppression of c-Maf in Treg cells

To investigate the role of Sox4 in chromatin regulation, we performed H3K27ac ChIP-sequencing, which detects active enhancer and promoter regions, on Sox4-overexpressing and control Treg cells. In Sox4-overexpressing cells, 3,868 genomic regions were more activated than in mock-infected control Treg cells (FDR < 0.1, depicted in blue and grey-blue), whereas 3,368 regions were repressed (FDR < 0.1, depicted in red and grey-red) (Figure 6A). The regions more active in control Treg cells included those around the *Il10* and *Ctla4*-gene loci, consistent with data of Sox4-mediated downregulated genes (Figure 1B). Among the regions more activated in Sox4-overexpressing Treg cells, DNA motifs recognized by Sox4 were most highly enriched, suggesting that Sox4 directly promotes chromatin activation at these sites (Figure 6B). In contrast, the regions more active in control Treg cells (i.e., those repressed by Sox4) lacked Sox4 motifs but were highly enriched for Maf recognition element (MARE), suggesting that Sox4 may suppress chromatin activity indirectly by inhibiting the Maf transcription factor rather than through direct binding (Figure 6B). Indeed, the regions more active in control Treg cells around the *Il10* and *Ctla4* loci substantially overlapped with previously reported c-Maf binding site in Th17 cells (Figure 6C)^22^.

**Figure 6.**
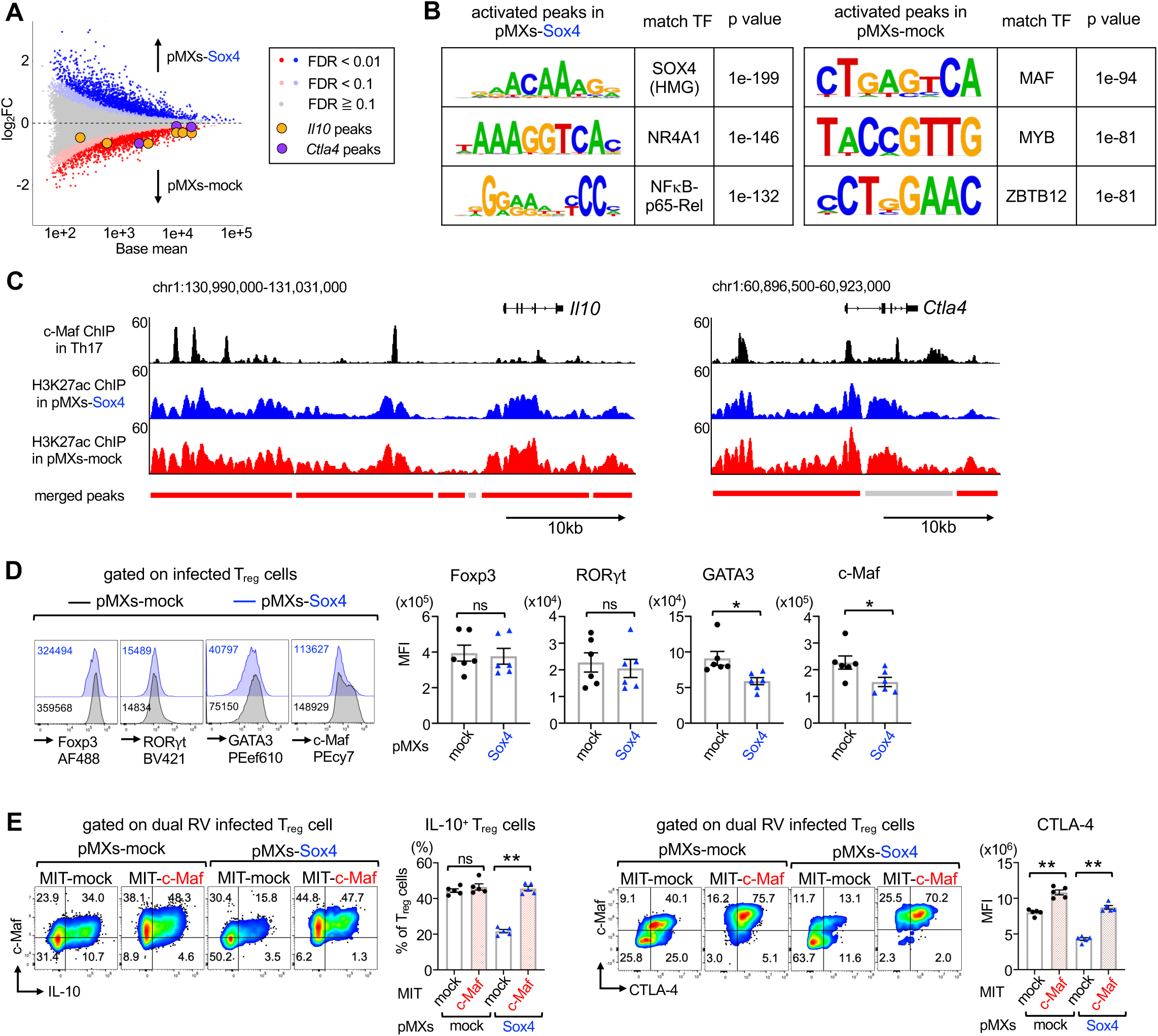
Sox4 decreases IL-10 and CTLA-4 production via the suppression of c-Maf in Treg cells. (**A**-**E**) Treg cells purified from Foxp3^YFP-cre^ mice were cultured in IL-10^+^Treg-polarizing conditions, infected with the indicated retroviruses, and analyzed 3 days after the infection. Infected retroviruses are either pMXs–IRES–NGFR (pMXs–mock) or pMXs-Sox4-IRES-GFP (pMXs-Sox4) (**A**-**D**), and either pMXs-mock or pMXs-Sox4 and either MSCV-IRES-Thy1.1 (MIT-mock) or MSCV-c-Maf-IRES-Thy1.1 (MIT-c-Maf) (**E**). (**A**) MA plot of tag counts on merged peaks derived from H3K27ac ChIP-seq. Significantly activated regions in pMX-mock-infected Treg cells (red) or pMX-Sox4-infected Treg cells (blue) are shown at FDR < 0.1. Similarly, less activated regions (FDR ≥ 0.1) are shown in gray, and intermediate regions (0.01 ≤ FDR < 0.1) in red-gray or blue-gray, respectively. Regions around *Il10* and *Ctla4* loci are surrounded by yellow and purple, respectively. (**B**) *de novo* motif analysis of the significantly activated regions in pMXs-mock- or pMXs-Sox4- infected Treg cells. (**C**) ChIP-seq tracks of H3K27ac in pMXs-mock- or pMXs-Sox4-infected Treg cells, and c-Maf in Th17 cells (GSE40918)^22^ around *Il10* and *Ctla4* loci are shown in red, blue, and black, respectively. The bottom bars indicate merged peaks, which are colored as in Figure 6A. (**D**) Representative histograms of indicated transcription factors in pMXs-mock- or pMXs-Sox4- infected Treg cells and the MFI of indicated transcription factors. n = 6, each. Unpaired t-test. *p<0.05, ns: not significant. (**E**) Representative flow cytometry profiles of IL-10 vs. c-Maf in retrovirus-infected Treg cells and the frequencies of IL-10-producing Treg cells or CTLA-4 MFI among infected Treg cells. n = 5, each. Two-way ANOVA followed by Tukey’s test. **p<0.01, ns: not significant.

Given these findings, we next explored whether Sox4 influences IL-10 and CTLA-4 expression by modulating the expression or activity of transcription factors such as c-Maf. Sox4 family proteins are known to regulate CD4□ T cell function through interactions with other transcription factors^19, 23^. Because c-Maf transcriptionally activates *Il10* and *Ctla4* gene loci^12, 24^, and GATA3 directly remodels the *Il10* locus in CD4□ T cells^12, 24, 25^, we hypothesized that Sox4 might suppress IL-10 production by downregulating these factors. Indeed, Sox4 overexpression in Treg cells reduced the protein levels of c-Maf and GATA3, but not those of Foxp3 and RORγt (Figure 6D). To test whether c-Maf can counteract Sox4-mediated repression, WT Treg cells were co-infected with MSCV-c-Maf-IRES-Thy1.1 (MIT-c-Maf) or control MSCV-IRES-Thy1.1 (MIT-mock) retrovirus together with pMXs-Sox4-IRES-NGFR (pMXs-Sox4) or control pMXs-IRES-NGFR (pMXs-mock) retrovirus. Analysis of doubly infected NGFR□Thy1.1□ cells revealed that c-Maf overexpression reversed the Sox4-induced suppression of IL-10 and CTLA-4 expression (Figure 6D), whereas GATA3 overexpression failed to rescue IL-10 suppression (Supplementary Figure 2). These results demonstrate that Sox4 suppresses the expression of IL-10 and CTLA-4 in Treg cells by repressing c-Maf-mediated transcriptional activation.

### Sox4 facilitates c-Maf degradation via the proteasome pathway through its 33 C-terminal residues

Since *Maf* was not identified as a Sox4-regulated gene in our RNA-seq analysis (Figure 1B), we examined c-Maf mRNA expression by quantitative RT-PCR in WT Treg cells upon Sox4 overexpression. Consistent with the RNA-seq results, c-Maf mRNA levels were not significantly affected by Sox4 overexpression (Figure 7A). The discrepancy between Sox4-mediated effects on c-Maf protein and mRNA expression suggests that Sox4 suppresses c-Maf protein levels at a post-transcriptional level.

**Figure 7.**
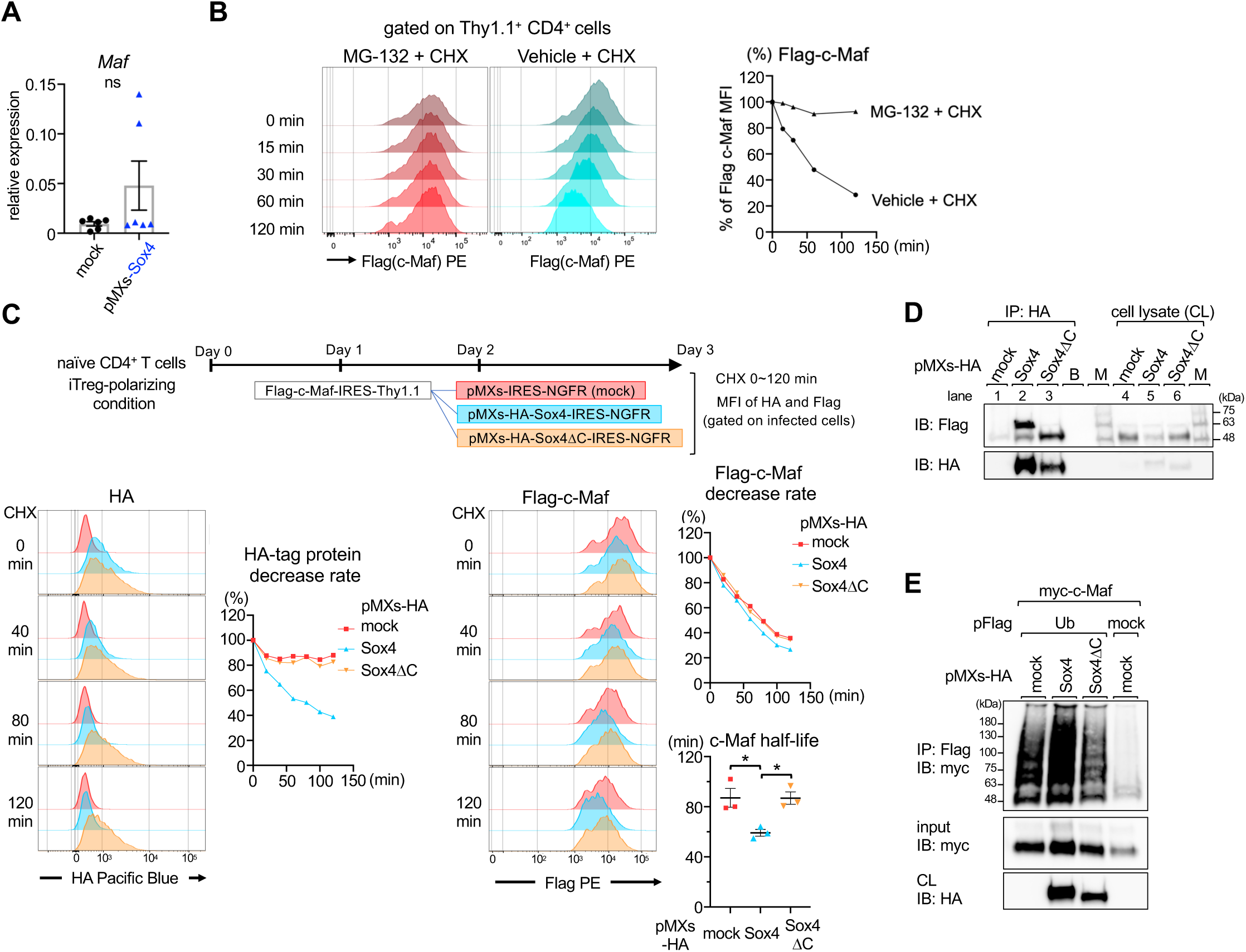
Sox4 physically interacts with c-Maf and facilitates its degradation via the ubiquitin-proteasome pathway through its C-terminal 33 residues. (**A**) WT Treg cells were cultured in IL-10^+^ Treg-polarizing conditions and infected with retroviruses of pMXs–mock or pMXs–Sox4. On day 3, infected cells were sorted, and the expression of Maf mRNA was analyzed by quantitative PCR. n = 6 each, unpaired t-test. ns: not significant. (**B**) Stability of c-Maf proteins in T cells. Naïve CD4^+^ T cells were stimulated and infected with Flag-c-Maf-IRES-Thy1.1 retrovirus on day 1. On day 3, cells were treated with either cycloheximide (CHX; 100 μg/mL) alone or CHX with MG-132 (20 μM) and analyzed at the indicated time points. Left panels: Representative histograms of Flag-Maf expression. Right panel: The decreased rate of Flag-c-Maf MFI in CHX-treated and CHX+MG-132-treated cells referred to the MFI of each untreated cell. (**C**) Effect of Sox4 on c-Maf stability. Upper panel: Schema of the assay. Naïve CD4^+^ T cells were cultured under iTreg-polarizing conditions and sequentially infected with Flag-c-Maf-IRES-Thy1.1 retrovirus on day 1 and either HA-Sox4-IRES-NGFR or HA-Sox4ΔC-IRES-NGFR retrovirus on day 2. On day 3, cells were treated with CHX and analyzed at the indicated time points. Lower left panels: Representative histograms of HA at the indicated time points and the decreased rate of the MFI of HA at the indicated time points referred to the MFI of each untreated cell. Lower right panels: Representative histograms of Flag-c-Maf at the indicated time points, the decreased rate of the MFI of Flag-c-Maf at the indicated time points referred to the MFI of each untreated cell, and the calculated half-life of Flag-c-Maf. n = 3 each. One-way ANOVA followed by Tukey’s test, *p < 0.05. (**D**) 293T cells were transfected with MSCV-Flag-c-Maf-IRES-Thy1.1 along with either pMXs-IRES-NGFR, pMXs-HA-Sox4-IRES-NGFR, or pMXs-HA-Sox4ΔC-IRES-NGFR. Two days after the transfection, cells were treated with MG-132 (10 μM) for 3 hours. Cell lysates were immunoprecipitated with anti-HA antibody and immunoblotted with anti-Flag antibody or anti-HA antibody. B: blank, M: molecular weight markers, CL: cell lysate. (**E**) 293T cells were transfected with Flag-Ub plasmids and myc-c-Maf plasmids along with either pMXs-IRES-NGFR, pMXs-HA-Sox4-IRES-NGFR, or pMXs-HA-Sox4ΔC-IRES-NGFR. Two days after the transfection, cells were treated with MG-132 (10 μM) for 3 hours, lysed in 6M guanidine buffer, and sonicated. c-Maf, which has natively 6x His sequences, was crudely purified by Cobalt-NTA Dynabeads, and imidazole-eluted protein complexes were subsequently immunoprecipitated with anti-Flag antibodies to purify ubiquitinated c-Maf. Imidazole-eluted protein complexes (input: middle panels) and anti-Flag immunoprecipitated protein complexes (upper panels) were immunoblotted with anti-Myc antibodies.

Next, we examined the stability of c-Maf proteins in T cells. In this experiment, CD4^+^ T cells were infected with Flag-c-Maf-IRES-Thy1.1 retrovirus, and the levels of Flag-c-Maf in Thy1.1^+^ cells were analyzed in the presence of cycloheximide (CHX), a protein synthesis inhibitor, with/without MG-132, a proteasome inhibitor (Figure 7B). In the presence of CHX, the levels of Flag–c-Maf rapidly decreased, and this decrease was completely blocked by MG-132, suggesting that c-Maf is rapidly degraded via the proteasome pathway in T cells.

We then investigated whether Sox4 accelerates the degradation of c-Maf. Given that the C-terminal 33 residues of Sox4 are essential for its own degradation^17^, we also examined the effect of a C-terminal 33 residue-deletion mutant of Sox4 (Sox4ΔC) on the degradation of c-Maf and Sox4. iTreg cells were sequentially infected with Flag-c-Maf-IRES-Thy1.1 retrovirus, followed by either HA-Sox4-IRES-NGFR, HA-Sox4ΔC-IRES-NGFR, or the control IRES-NGFR retroviruses. The MFI of Flag and HA in Thy1.1^+^NGFR^+^ iTreg cells was analyzed in the presence of CHX (Figure 7C). Consistent with previous reports, Sox4 was quickly degraded in iTreg cells compared to Sox4ΔC (Figure 7C, lower left panels). Despite this higher instability of Sox4 in iTreg cells, the calculated half-life of c-Maf co-expressed with Sox4 was significantly lower than that with Sox4ΔC (Figure 7C, lower right panels). These results indicate that the C-terminal region of Sox4 is essential for both Sox4-mediated c-Maf degradation and degradation of Sox4 itself.

### Sox4 physically interacts with c-Maf and facilitates its degradation via the ubiquitin proteasome pathway

We next investigated the physical interaction between Sox4 and c-Maf. In this experiment, MG-132 was added to prevent the rapid degradation of Sox4 and c-Maf. Our co-immunoprecipitation experiments revealed that both HA-Sox4 and HA-Sox4ΔC interacted with Flag-c-Maf (Figure 7D). Notably, the Flag-c-Maf co-immunoprecipitated with HA-Sox4 exhibited two distinct bands: a minor band at 52 kDa and a major band at 61 kDa (Figure 7D, upper panel, lane 2). The 61 kDa band was approximately 9 kDa larger than the Flag-c-Maf observed in cell lysates, suggesting that this form likely represents monoubiquitinated Flag-c-Maf. Consistent with the results of Figure 6C, we observed a decrease in Flag-c-Maf levels when co-expressed with HA–Sox4 but not with HA-Sox4ΔC in total cell lysates (Figure 7D, upper panel, lanes 5 and 6). This observation further supports the notion that Sox4 but not Sox4ΔC promotes the degradation of c-Maf. These results indicate that the C-terminal domain of Sox4 is dispensable for its interaction with c-Maf but is essential for mediating c-Maf degradation, suggesting a two-step mechanism where Sox4 first binds to c-Maf and subsequently facilitates its degradation through a process that requires the C-terminal domain.

Finally, we examined whether Sox4 enhances the ubiquitination of c-Maf (Figure 7E). 293T cells were transfected with myc-c-Maf along with either pFlag-Ub or pFlag-mock plasmids and either HA-Sox4, HA-Sox4DC, or mock plasmids. Because c-Maf naturally contains 6x His sequences, ubiquitinated c-Maf was crudely purified using Cobalt–NTA beads under denaturing conditions and then subjected to Flag immunoprecipitation to pull down Flag-polyubiquitinated c-Maf proteins. This analysis revealed that HA-Sox4 significantly enhanced the polyubiquitination of myc–c-Maf, as evidenced by a pronounced smear pattern in the Western blot analyses (Figure 7E, upper panel, lane 2). This enhancement was notably more substantial than the effects observed with HA-Sox4ΔC or mock controls (Figure 7E, upper panel, lane 3 and lane 1, respectively). As anticipated, in the absence of pFlag–Ub, no polyubiquitination smear was detected (Figure 7E, upper panel, lane 4). This observation serves as a crucial negative control, confirming the specificity of our ubiquitination assay. Taken together, Sox4 physically interacts with c-Maf and enhances its degradation by the ubiquitin-proteasome pathway through a process that requires the C-terminal domain of Sox4.

## DISCUSSION

This study identified Sox4 as a novel regulator of IL-10 production in Treg cells, which is crucial in controlling allergic airway inflammation. Our findings reveal that Sox4 is a negative regulator of IL-10 and CTLA-4 expression in Treg cells, influencing their suppressive function on Th2 cell proliferation. Collectively, these data establish Sox4 as a key molecular brake on Treg-mediated suppression in type 2 inflammation. Mechanistically, we found that Sox4 suppresses c-Maf protein levels, with c-Maf degradation mediated by Sox4’s C-terminal domain via the proteasome pathway. This mechanistic link provides a direct explanation for the observed reduction in IL-10 production in Sox4-expressing Tregs.

To explore IL-10 regulation specific to Treg cells, RNA-seq analysis of IL-10^+^ Treg, IL-10^−^ Treg, and IL-10^+^ Tconv (VitD3+Dex) revealed a decrease in *Sox4* and an increase in *Rorc* and *Arnt2* expression in IL-10^+^ Treg cells (Figure 1A). *Rorc*, the master regulator of Th17 cells, is typically suppressed by IL-2. Given that c-Maf, expressed in IL-10-producing T cells, suppresses IL-2, it is plausible that *Rorc* upregulation might occur through c-Maf-mediated IL-2 suppression^12^. Arnt2 (aryl hydrocarbon receptor nuclear translocator 2), known to heterodimerize with Ahr (aryl hydrocarbon receptor)^26^, influences the reciprocal differentiation of Treg and Th17 cells^27^. Thus, Arnt2 may represent an additional regulatory node linking Sox4 signaling to the Treg/Th17 balance.

We identified fifteen Sox4-mediated upregulated genes and thirteen downregulated genes in Treg cells (Figure 1B). To elucidate how Sox4 modulates Treg function, we focused on genes likely to affect their suppressive capacity. Among these, *Igfbp4*, *Il10*, and *Ctla4* emerged as notable candidates. Although IGFBP-4 (insulin-like growth factor binding protein 4) has been reported to suppress Treg proliferation by inhibiting IGF (insulin-like growth factor) function^28^, Sox4-cKO Treg cells did not exhibit increased proliferation, suggesting the Sox4-IGFBP4 axis may be less critical. In contrast, CTLA-4 expression was markedly reduced by Sox4 both in vitro and in vivo (Figure 2A and 5). Because CTLA-4 expressed on Treg cells mediates trans-endocytosis of CD80/86 on antigen-presenting cells^29, 30^, future studies should determine whether Sox4 influences this process.

Sox4 deficiency in Treg cells ameliorated eosinophilic inflammation in the airway (Figure 3), contrary to results in T cell-specific Sox4-deficient mice^19^. This apparent discrepancy likely reflects the dual, context-dependent roles of Sox4 in Th2 and Treg lineages. Sox4 suppresses Th2 responses by suppressing GATA3 binding to *Il5* promoter without inhibiting GATA3 mRNA expression^19^. Therefore, in T cell–wide Sox4 deletion, elevated IL-5 from Sox4-deficient Th2 cells may override the anti-inflammatory effects of Sox4-deficient Tregs. With respect to the relationship between GATA3 and Sox4, we observed Sox4-dependent inhibition of GATA3 both in vitro and in vivo in IL-10-producing Treg cells (Figures 5 and 6D). Because Sox4 induces the degradation of GATA3 via ubiquitin E3 ligase Fbw7 in breast cancer cells^31^, and Sox12, another SoxC protein, similarly promotes Fbw7-mediated GATA3 degradation in Th2 cells^23^, it is likely that Sox4 similarly degrades GATA3 in IL-10-producing Treg cells.

Neutralizing IL-10R before allergen challenge eliminated the protective activity of Sox4-deficient Treg cells against type 2 airway inflammation (Figure 4), while IL-10R blockade during sensitization or throughout challenge had no significant effect (data not shown). During the sensitization phase, IL-10 produced predominantly by B cells promotes CCL20 expression in airway epithelium, recruits dendritic cells, and facilitates allergen priming. Accordingly, IL-10 blockade during this window dampens allergic sensitization and asthma development^32^. In later stages, however, IL-10 signaling is indispensable for Treg cell–mediated control of allergic inflammation^33^. Mechanistically, IL-10 upregulates granzyme B in Th2 cells, limiting their persistence and restraining airway inflammation^10^. Thus, the impact of IL-10 is spatially and temporally segregated, with local lung IL-10 production determining its functional outcome. Within this context, Sox4-deficient Treg cells in the lung upregulate IL-10 and CTLA-4 and effectively suppress Th2-driven inflammation.

H3K27ac ChIP-seq analysis showed that not only Sox4 but also Nr4a1 and Rel binding motifs were enriched in the activated regions of Sox4-overexpressing Treg cells (Figure 6B). Nr4a family members and Nfkb family members are essential for thymic Treg differentiation by promoting Foxp3 expression and establishing Treg lineage stability^34, 35^. However, thymic Treg differentiation was comparable between Sox4-cKO and Sox4-cHet mice (data not shown). Since Foxp3-Cre-mediated recombination occurs after Foxp3 expression is initiated in CD4 single-positive thymocytes^36^, Sox4 deletion in Foxp3^YFP-cre^ Sox4^f/f^ (Sox4-cKO) mice would occur downstream of the critical early stages of thymic Treg commitment^37^, potentially masking earlier requirements for Sox4. Therefore, studies using CD4^cre^ Sox4^fl/fl^ mice, in which gene deletion precedes Foxp3 induction, are needed to clarify whether a Sox4–Nr4a or Sox4–Nfkb axis plays a role in thymic Treg development.

H3K27ac ChIP-seq analysis further demonstrated that Sox4 represses c-Maf– and Myb-dependent active enhancer and promoter regions. Myb is indispensable for effector Treg differentiation, and its deficiency leads to impaired proliferation and differentiation, culminating in multi-organ inflammation^38^. Myb controls key effector Treg genes, including *Gata3*, *Maf,* and *Ctla4*. However, because IL-10 is not a known Myb target and deposit ChIP-seq data of Myb showed no binding around the Il10 loci (data not shown), the Sox4–Myb axis likely contributes less to Sox4-mediated IL-10 suppression in Treg cells than the Sox4–c-Maf pathway.

Finally, we confirmed that Sox4 suppresses IL-10 production by suppressing c-Maf protein in Treg cells (Figure 6D) and that the C-terminal 33 residues of Sox4 induce the degradation of c-Maf (Figure 7). It has been shown that c-Maf interacts with several ubiquitin E3 ligases, including RNF113, HERC4, UBR5, and HUWE1^39^. Therefore, Sox4 may enhance the degradation of c-Maf by making a ternary complex with these ubiquitin E3 ligases. Further studies are required to identify the specific E3 ligase responsible for Sox4-induced ubiquitination of c-Maf.

In conclusion, we have uncovered a novel mechanism by which Sox4 induces c-Maf degradation, thereby suppressing IL-10 production in Treg cells. Sox4-deficient Treg cells exhibit enhanced IL-10 production via c-Maf upregulation and demonstrate improved efficacy in ameliorating eosinophilic inflammation in a murine asthma model. These findings offer new insights into regulating IL-10-producing Treg cells and provide strategies for controlling type 2 inflammation.

## ACKNOWLEDGEMENTS

Ms. M. Yoshino for technical help. This work was supported by a Health Labor Sciences Research Grant on Allergic Disease and Immunology from the Ministry of Health, Labor, and Welfare of Japan, JST (Moonshot R&D) (Grant Number JPMJMS2025), AMED under Grant Number JP223fa627003, and the Innovative Medicine CHIBA Doctoral WISE Program at Chiba University from the Ministry of Education, Culture, Sports, Science, and Technology. The authors have no conflicting financial interests.

## SUPPLEMENTARY INFORMATION

Supplementary Figures 1 and 2.

## AUTHOR CONTRIBUTIONS

Study conception and design, A.S. Y.H. and K.Suga.; Acquisition of data, Y.H., K.Suga, A.I., and A.S.; Provide important materials, M.Y. and V.L., Analysis and interpretation of data, K.A., T.K., T.I., J.I, A.I., S.T., K Suzuki., and H.N.; Wrote the paper, A.S. K.Suga., Y.H., A.I., and H.N.

## STAR⍰METHODS

Detailed methods are provided in the online version of this paper and include the following:

### RESOURCE AVAILABILITY

#### Lead contact

Further information and requests for resources and reagents should be directed to and will be fulfilled by the lead contact, Akira Suto.

#### Materials availability

Materials used in this study are available from the lead contact upon request.

#### Data and code availability

The datasets used to generate figures from the present study have not been deposited in a public repository but are available on request from the lead contact.

This paper does not report the original code.

Any additional information required to reanalyze the data reported in this paper is available from the lead contact upon request.

### EXPERIMENTAL MODEL AND STUDY PARTICIPANT DETAILS

#### Mice

C57BL/6 mice were purchased from Charles River Laboratories (Kanagawa, Japan). C57BL/6 background Sox4^f/f^ mice generated by Dr. Lefebvre^40^ are kindly provided by Dr. Yamashita. Foxp3^YFP-cre^ mice were purchased from Jackson Laboratory. Sox4^f/f^ mice were crossed to Foxp3^YFP-cre^ mice. Because Foxp3^YFP-cre^ mice have been shown to induce ectopic germline recombination, we employed a quantitative PCR-based genotyping method to investigate possible ectopic recombination outside the Treg lineage. We exclude mice with > 30% ectopic recombination that harbor a germline Sox4 recombination allele. All mice were housed in microisolator cages under specific pathogen-free conditions. The Chiba University Animal Care and Use Committee approved protocols for animal experiments.

### METHOD DETAILS

#### Re-analysis of public RNA-sequencing data

RNA-sequencing data were obtained through Gene Expression Omnibus (accession number GSE106464). After adapter removal by TrimGalore, raw sequence data were aligned to the mm10 genome by HISAT2, and read counts were generated using the featureCounts program. After the normalization of raw count values, differences in transcript expression between groups were assessed using the DESeq2 package in R.

#### Analysis of Sox4-regulated genes in Treg cells

Treg cells were isolated from the spleen and lymph nodes of Foxp3^YFP-cre/YFP-cre^ mice and Foxp3^YFP-cre/YFP-cre^ Sox4^f/f^ mice (WT Treg cells and Sox4-cKO Treg cells, respectively, n = 2 each). In another set of experiments, sorted WT Treg cells were infected with Sox4 retrovirus (pMXs-Sox4) or a control retrovirus (pMXs-empty), and the infected cells were isolated by the expression of NGFR (n = 2, each) (Figure 1B). Both pairs of cells were stimulated with PMA and ionomycin (P+I) and subjected to RNA sequencing. RNA-seq libraries were prepared using a QuantSeq 3’ mRNA-Seq Library Prep Kit (Lexogen, Greenland, NH). Sequencing was performed on an Illumina NextSeq500 using a NextSeq 500/550 High Output Kit v2.5 (Illumina, San Diego, CA) in a 75-base single-read mode. RNA-seq reads were mapped with STAR v2.7.6a, and count data were calculated with HTSeq-count v1.99.2. A web-based integrative transcriptome analysis RNAseqChef package was used to identify DEGs and to normalize count data. GEO accession number is GSE282323. Genes upregulated by overexpressing Sox4 and downregulated by Sox4-cKO were considered genes upregulated by Sox4. Genes downregulated by Sox4 are vice versa.

#### Intracellular staining of cytokines and transcription factors and flow cytometry analysis

To detect intracellular cytokines, cells were stimulated with PMA and ionomycin for 4 hours in the presence of BD GolgiPlug (BD Biosciences, Franklin Lakes, NJ). Intracellular cytokine and transcription factor staining were performed using an eBioscience Foxp3 transcription factor fixation/permeabilization kit (Thermo Fisher Scientific, Waltham, MA), as described previously^41^. Cells were analyzed by Novocyte Penteon (Agilent Technologies, Santa Clara, CA) or BD FACSCanto II. Flow cytometry profiles were analyzed using the FlowJo software ver. 10.9 (BD Biosciences).

#### Plasmids and retrovirus-mediated gene expression

pMXs-IRES-NGFR is a retrovirus vector expressing human NGFR under the regulation of an internal ribosome entry site (IRES). pMXs-Sox4-IRES-NGFR is a kind gift from Dr. M. Yamashita (Ehime University, Ehime, Japan). HA-tag was fused to Sox4 by PCR amplification, and HA-Sox4 was subcloned into pMXs-IRES-NGFR to create pMXs-HA-Sox4-IRES-NGFR. Sox4 mutants lacking the C-terminal 33 residues (Sox4ΔC) were generated using a KOD-Plus-mutagenesis kit according to the manufacturer’s instructions. MSCV-c-Maf-IRES-Thy1.1 and pFlag-Ub are previously described^23, 42^. Flag or myc tag was fused to c-Maf by PCR amplification, and Flag–c-Maf or myc–c-Maf was subcloned into MSCV-IRES-Thy1.1 to create MSCV-Flag-c-Maf-IRES-Thy1.1 or MSCV-myc-c-Maf-IRES-Thy1.1, respectively. All constructs were verified by sequencing. Retrovirus-mediated gene induction for T cells was performed by a RetroNectin-bound virus infection method (Takara Bio, Otsu, Japan) ^43^.

#### Cell isolation and cell culture

Naïve CD4^+^ T cells were isolated from lymph nodes or spleen using a MojoSort mouse naïve T cell isolation kit (Biolegend, San Diego, CA) according to the manufacturer’s instructions. CD45.1^+^ CD62L^+^ naïve CD4^+^ T cells or Foxp3^YFP+^ CD25^+^ Treg cells were purified using FACSMelody cell sorter (BD, Franklin Lakes, NJ). For the development of IL-10-producing Treg cells, Treg cells were stimulated with plate-bound anti-CD3ε mAb (1 μg/ml) in the presence of anti-CD28 mAb (1 μg/ml) under IL-10^+^Treg-polarizing conditions (IL-4 (10 ng/ml) and IL-2 (10 ng/ml)) as previously reported^13^. For development of iTreg cells, naïve CD4 T cells were stimulated with plate-bound anti-CD3ε mAb (1 μg/ml) in the presence of anti-CD28 mAb (1 μg/ml) under iTreg-polarizing conditions TGF-β (3 ng/ml), IL-2 (10 ng/ml), anti-IL-4 mAb (10 μg/ml), and anti-IFN-γ mAb (10 μg/ml).

#### Suppression assay

Treg cells were sorted from the spleen and lymph nodes of Foxp3^YFP-Cre^ mice by a FACSMelody cell sorter. Naïve CD45.1^+^ CD4^+^ responder T cells were purified from CD45.1 congenic mice and stained with CellTrace Violet according to the manufacturer’s instructions (Thermo Fisher Scientific). Suppression assay was performed as described previously^18^. Where indicated, anti-CTLA-4 antibodies (100 μg/ml) were added to the culture medium.

#### Transwell experiments

Transwell experiments were performed in 96-well plates with a pore size of 0.4 μM (Millipore). Naïve CD4^+^ T cells (5 x 10^4^ cells) were stimulated with dynabeads mouse T-activator CD3/CD28 (Thermo Fisher Scientific) (anti-CD3/CD28 beads) (1 x 10^5^ beads) in bottom chamber, and naïve CD4 T cells (2.5 x 10^4^ cells) and WT Treg cells (5 x 10^3^ cells) were stimulated with anti-CD3/CD28 beads (5 x 10^4^ beads) in upper chamber in the presence of IL-4 (10 ng/ml) and anti-IFN-γ antibody (10 μg/ml). Where indicated, anti-IL-10 receptor (IL-10R) antibodies (10 μg/ml) were added to the culture medium. The numbers of GATA3^+^ Th2 cells in the bottom chambers were analyzed on day 3.

#### HDM-induced allergic airway inflammation

Mice were sensitized and challenged with intratracheal administration of HDM extracts (Greer Laboratories) as described previously^23^. In brief, mice were sensitized intratracheally with HDM (1 μg in 25 μl PBS) at day 0 and day 7 and were challenged with HDM (1 μg in 25 μl PBS) at day 18. Forty-eight hours after the HDM challenge, asthmatic responses were evaluated. Single-cell suspensions of total lung cells were obtained using gentleMACS^TM^ Octo Dissociator (Miltenyi Biotec). Eosinophils were defined as CD45^+^ CD11b^+^ CD11c^-^ Siglec-F^+^ cells, CD4^+^ T cells as CD45^+^ CD11b^-^ CD4^+^ CD3^+^ cells. Lung cells were stimulated with PMA plus ionomycin (PMA+I) in the presence of brefeldin A for 4 h, and the production of IL-5, IL-10, and IL-13 was evaluated by intracellular cytokine staining. Transcription factors and CTLA-4 were analyzed without stimulation. Lung tissues were fixed in 4% paraformaldehyde, embedded in paraffin, and sectioned at 4μm, followed by staining with hematoxylin and eosin and periodic acid-Schiff (PAS). Lung inflammation and goblet cell hyperplasia were graded as previously described^44^. Briefly, inflammatory-cell numbers were graded as follows: 0, normal; 1, few cells; 2, a ring of inflammatory cells that was 1 cell-layer deep; 3, a ring of inflammatory cells that was 2–4 cell-layers deep; and 4, a ring of inflammatory cells that was >4 cell-layers deep. The goblet-cell numbers were graded as follows: 0, <0.5% PAS-positive cells; 1, <25%; 2, 25–50%; 3, 50–75%; and 4, >75%. Where indicated, anti-IL-10R antibodies (250μg) were intraperitoneally administered 6h before the challenge.

#### Mixed bone marrow chimeras (mBMC)

Bone marrow cells obtained from Foxp3^YFP-cre^ mice (CD45.1^+^ background) and CD45.2^+^ background Foxp3^YFP-cre^ Sox4^f/f^ mice (2 x 10^6^ cells, each) were mixed and injected intravenously into CD45.1^+^/CD45.2^+^ background C57BL/6 mice after total body irradiation (9.5 Gy). mBMC mice were used 8 weeks after the bone marrow reconstitution.

#### ChIP-seq analysis

Treg cells were sorted from the spleen and lymph nodes of Foxp3^YFP-Cre^ mice and infected with pMXs-IN-HASox4 or pMXs-IN-mock and cultured in IL-10^+^Treg-polarizing conditions for three days. ChIP-seq was performed as previously described with minor modifications^45^. 1 x 10^5^ NGFR□ infected Treg cells were sorted and crosslinked with 1% formaldehyde for 5 minutes at RT. Chromatin DNA was extracted and sonicated using a PicoRuptor (Diagenode). Immunoprecipitation was performed with an anti-H3K27ac antibody (Abcam, ab4729) and Protein A dynabeads (Thermo Fisher Scientific). Sequencing libraries were prepared using the NEBNext Ultra II DNA Library Prep Kit, NEBNext Multiplex Oligos (New England BioLabs). The resulting libraries were converted for DNBSEQ sequencing using the MGIEasy Universal Library Conversion Kit (MGI Tech) and sequenced on a DNBSEQ-G400RS platform in 100 bp paired-end mode. Raw data of c-Maf ChIP-seq in Th17 cells (GSE40918) were reanalyzed. Adapter sequences were trimmed using Trimmomatic v0.39^46^, and the filtered reads were aligned to the mouse reference genome (mm10) using Bowtie2^47^. Visualization and peak calling were performed using HOMER v5.1^48^. Shared peaks between two replicates of both cell types were merged, and tag counts were calculated on the merged peaks using annotatePeaks.pl of HOMER. Differentially activated peaks were calculated by DESeq2^49^, and an MA plot was generated by R. *de novo* motif analysis was performed using findMotifsGenome.pl of HOMER.

#### Coimmunoprecipitation (co-IP) analysis

293T cells were transfected with the indicated vectors using an Effectene Transfection Reagent (Qiagen). Thirty-six hours after the transfection, cells were lysed in IP lysis buffer [50 mM NaCl, 50 mM Tris, 0.5% Nonidet P-40, 10 mM sodium pyrophosphate, 5 mM EDTA, 50 mM NaF, and 1% protease inhibitor cocktail (Merk, St. Louis, MO, USA)]. Whole-cell extracts were immunoprecipitated with anti-Flag-M2 antibodies (Merk) conjugated with Protein G dynabeads (Thermo Fisher Scientific).

#### Ubiquitination assays

293T cells were transfected with MSCV-myc-c-Maf-IRES-Thy1.1 and Flag-tagged Ub (Flag-Ub) along with either pMXs-IRES-NGFR or pMXs-HA-Sox4-IRES-NGFR or pMXs-HA-Sox4ΔC-IRES-NGFR. Two days after the transfection, cells were treated with MG-132 (10 μM) for 3 hr. Ten % of transduced cells were lysed in IP lysis buffer and used as an input sample. Residual 90 % of transduced cells were lysed in 1 ml of buffer A [6 M guanidinium-HCl, 0.1 M sodium phosphate (pH 8.0), and 10 mM imidazole] ^50^, sonicated for 10 seconds 3 times, and mixed with 50 μl of Cobalt-NTA Dynabeads (ThermoFisher Scientific) on a rotator for 30 min at room temperature. Cobalt–NTA Dynabeads were sequentially washed with 1 ml of buffer A, with 1 ml of buffer A diluted in binding/wash buffer at 1:4, and with 1 ml of binding/wash buffer [50 mM sodium phosphate (pH 8.0), 300mM NaCl, and 0.02 % Tween]. Because c-Maf naturally contains 6x His sequences that bind Cobalt–NTA Dynabeads, crudely purified myc–c-Maf proteins were eluted by boiling the Cobalt–NTA Dynabeads in 2x sample buffer supplemented with 300 mM imidazole. The same amounts of myc-c-Maf proteins were diluted in PBS to adjust the SDS concentration to 0.1 %. Ten % of diluted solutions were left as a control sample, and 90 % of diluted solutions were mixed with 20 μl of Protein G dynabeads coated with anti-Flag M2 antibodies (Sigma) on a rotator for 30 min at room temperature to purify ubiquitinated myc–c-Maf proteins. After beads were washed 3 times with binding/wash buffer, ubiquitinated myc–c-Maf proteins were eluted by boiling in 2x sample buffer and analyzed by immunoblotting with HRP-conjugated anti-Myc antibody.

### QUANTIFICATION AND STATISTICAL ANALYSIS

Data are summarized as means ± SEM. The statistical analyses of the results were performed using the unpaired Student’s t-test, one-way ANOVA followed by Tukey’s test, or two-way ANOVA followed by Tukey’s test. P<0.05 was considered significant.

### KEY RESOURCES TABLE

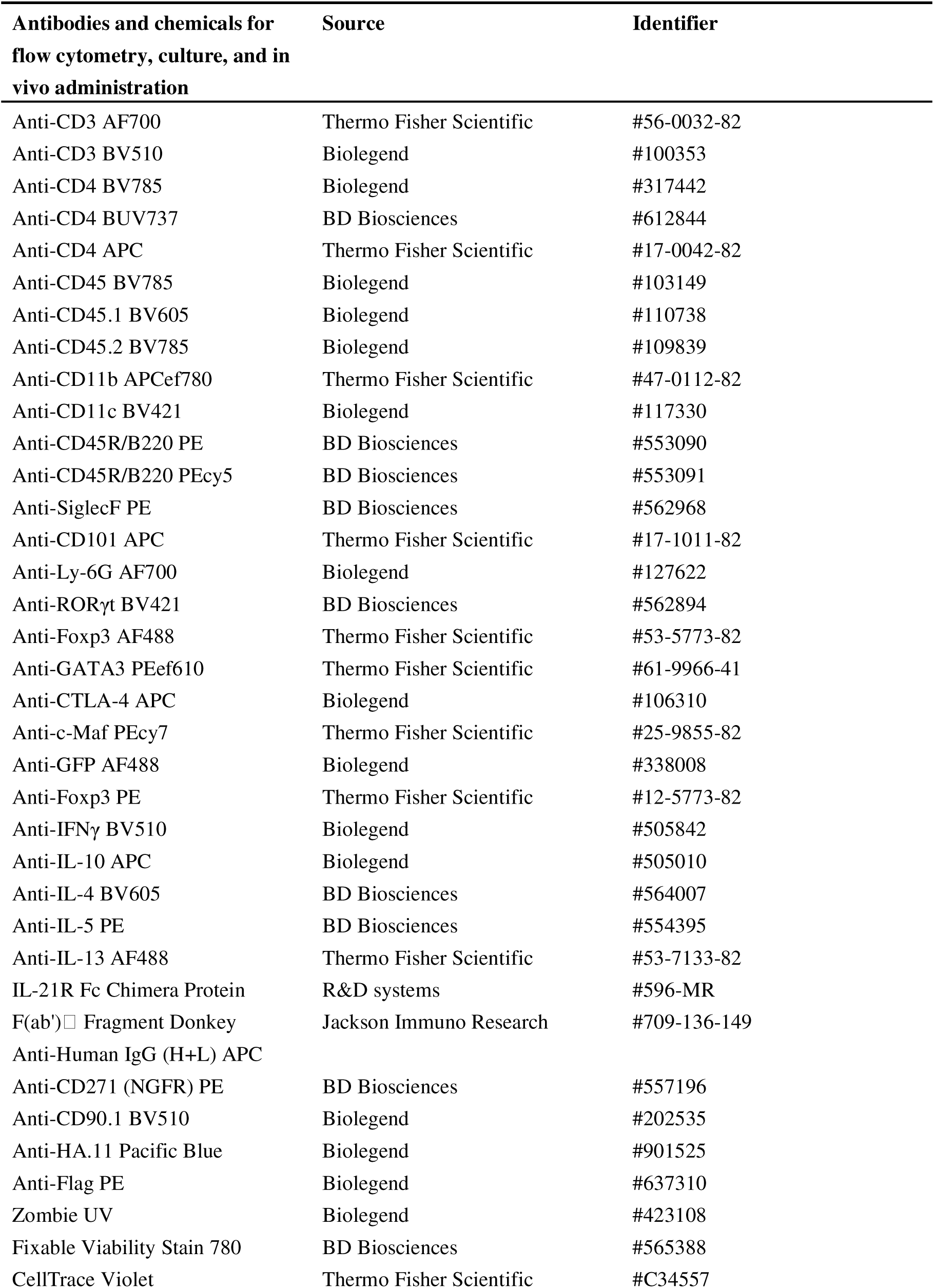
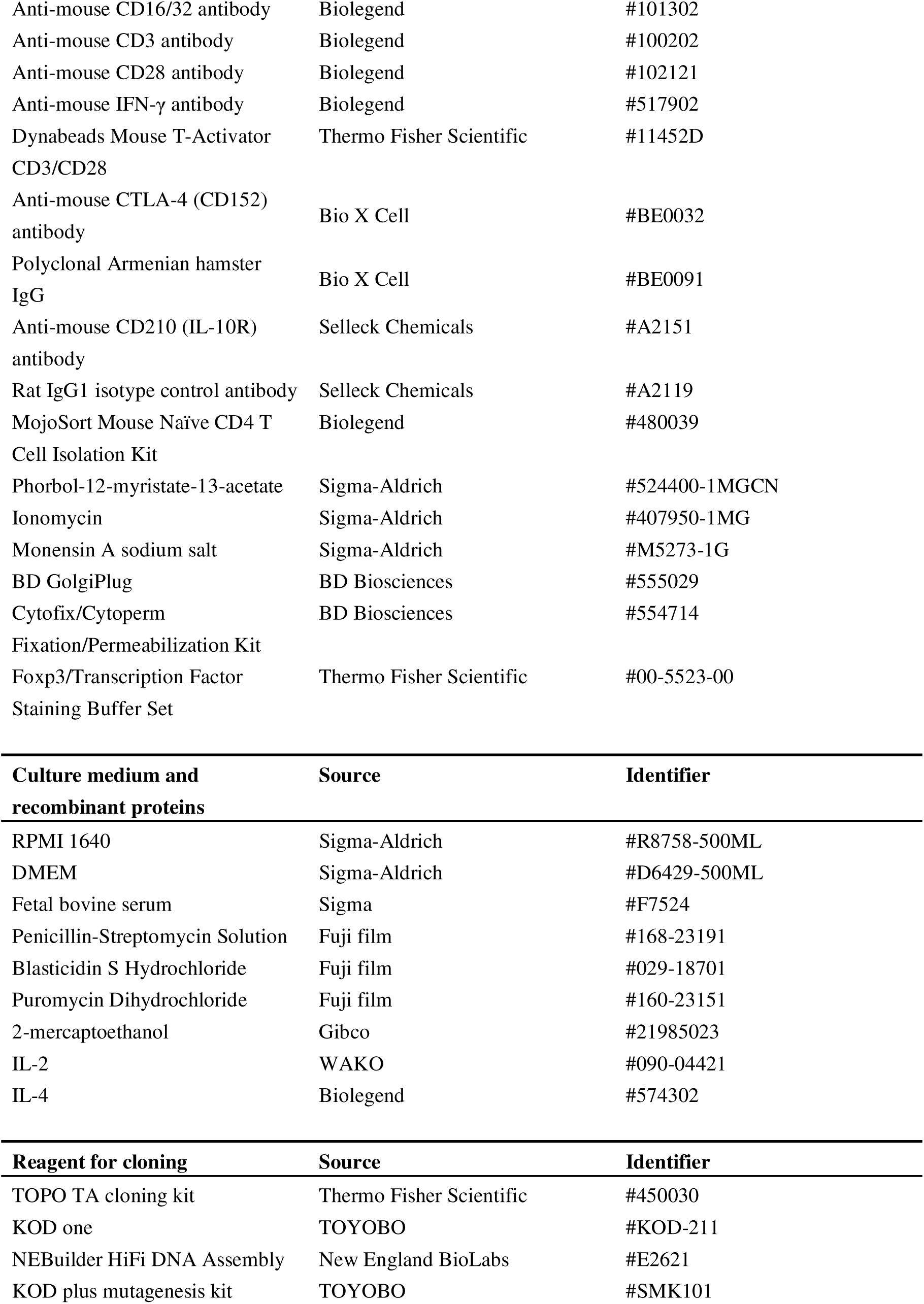
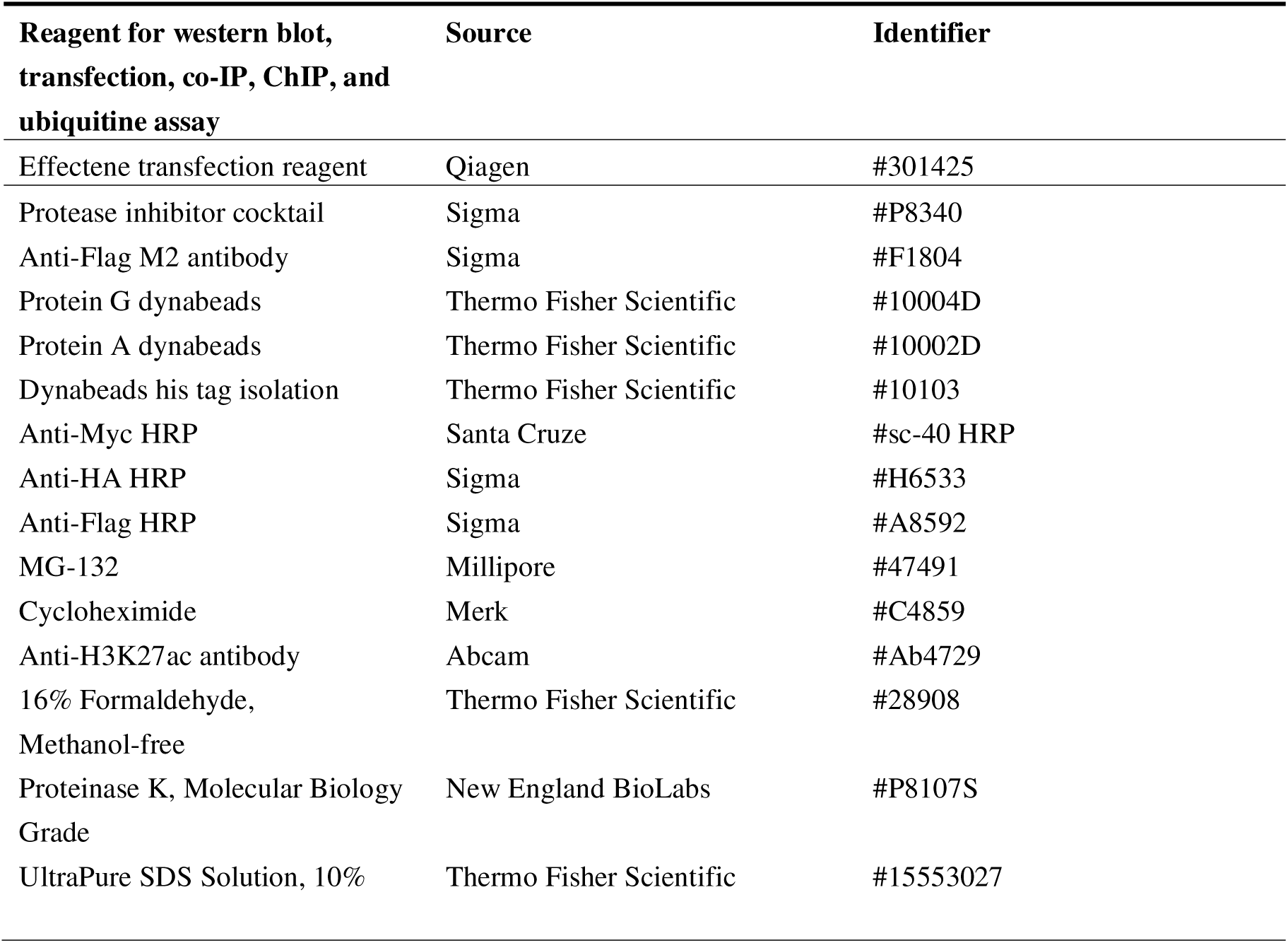

## SUPPLEMENTAL FIGURE LEGENDS

**Supplementary Figure 1. Sox4-cKO Treg cells show no significant difference in the expression of CTLA-4 and IL-10 in mediastinal lymph nodes under type 2 inflammation**

(**A, B**) Foxp3^YFP-cre/wild-type^ Sox4^f/f^ female mice were sensitized and challenged, and the draining mediastinal lymph nodes were analyzed as described in Figure 5A and B. One-way ANOVA followed by Tukey’s test. n = 6 each. ns: not significant.

(**C, D**) The mBMC mice described earlier were sensitized and challenged, and the draining mediastinal lymph nodes were analyzed as described in Figure 5C and D. n = 8 for HDM and n = 6 for PBS. One-way ANOVA followed by Tukey’s test. ns: not significant.

**Supplementary Figure 2. Overexpression of GATA3 fails to rescue Sox4-mediated suppression of IL-10 expression.**

Treg cells purified from Foxp3^YFP-cre^ mice were cultured, infected with retroviruses, and analyzed as described in Figure 6D. Infected retroviruses are either pMXs-mock or pMXs-Sox4 and either MSCV-IRES-Thy1.1 (MIT-mock) or MSCV-GATA3-IRES-Thy1.1 (MIT-GATA3). n = 5, each. Two-way ANOVA followed by Tukey’s test. ns: not significant.

