## Supplementary figures for "Sox4 in Treg Cells Suppresses IL-10 Production via c-Maf Degradation and Exacerbates Type 2 Inflammation"

Fig.S1 **Sox4-cKO Treg cells show no significant difference in the expression of IL-10 and CTLA-4 in mediastinal lymph nodes under type 2 inflammation**

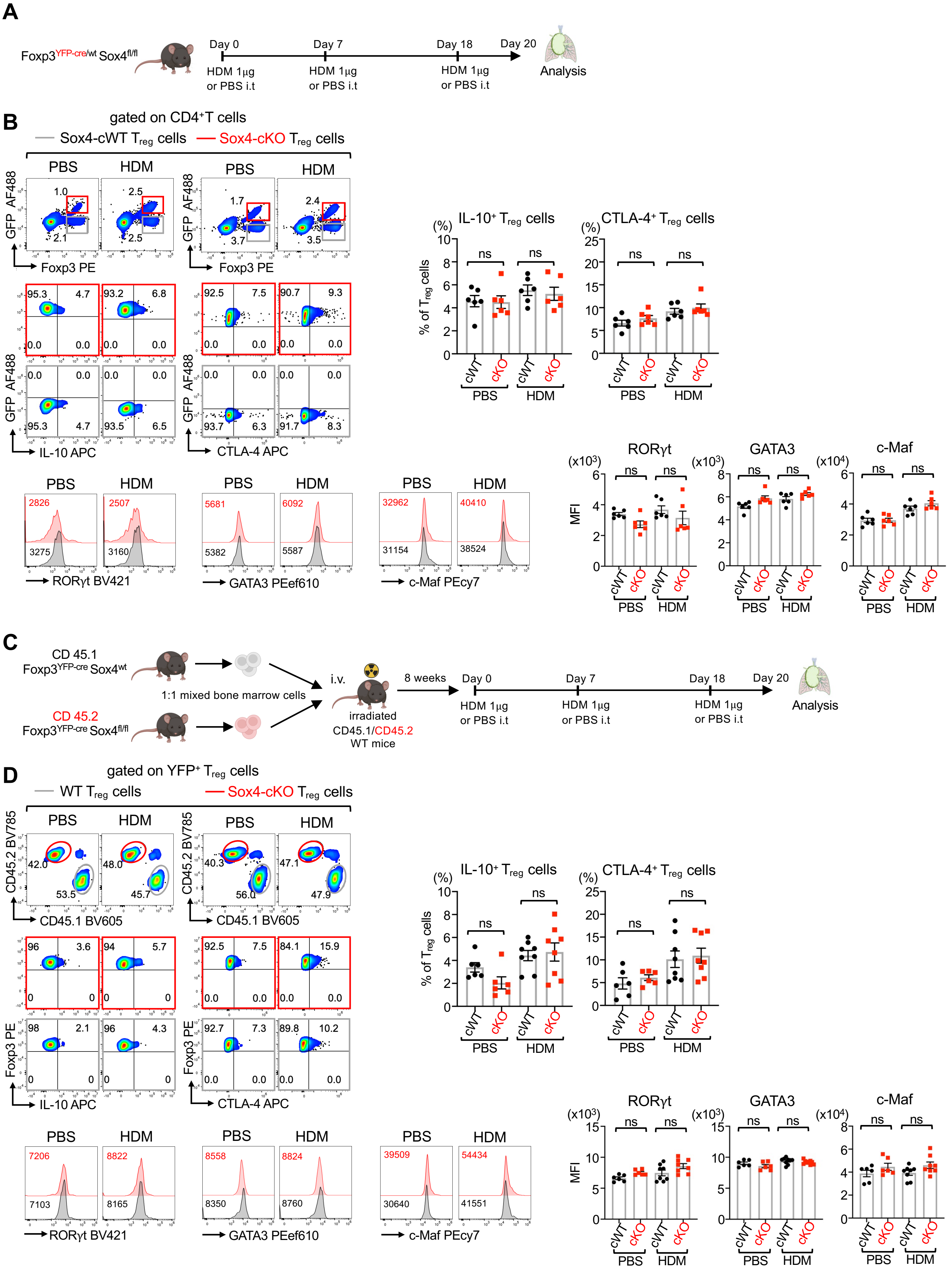

Fig.S2 **Overexpression of GATA3 fails to rescue Sox4-mediated suppression of IL-10 expression**

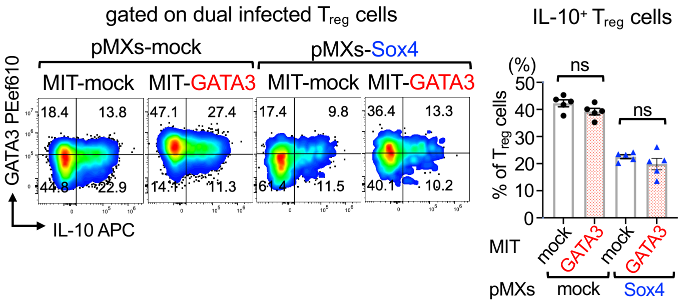
